# The role of macrophage scavenger receptor 1 (Msr1) in prion pathogenesis

**DOI:** 10.1101/2020.11.17.386433

**Authors:** Bei Li, Meiling Chen, Adriano Aguzzi, Caihong Zhu

## Abstract

The progression of prion diseases is accompanied by the accumulation of prions in the brain. Ablation of microglia enhances prion accumulation and accelerates disease progression, suggesting that microglia play a neuroprotective role by clearing prions. However, the mechanisms underlying the phagocytosis and clearance of prion are largely unknown. The macrophage scavenger receptor 1 (Msr1) is an important phagocytic receptor expressed by microglia in the brain, and is involved in the uptake and clearance of soluble amyloid-β. We therefore asked whether Msr1 might play a role in prion clearance, and assessed the scavenger function of Msr1 in prion pathogenesis. We found that Msr1 expression was upregulated in prion-infected mouse brains. However, Msr1 deficiency did not change prion disease progression or lesion patterns. Prion deposition in Msr1 deficient mice was similar to their wild type littermates. In addition, prion-induced neuroinflammation was not affected by Msr1 ablation. We conclude that Msr1 does not play a major role in prion pathogenesis.

## Introduction

Prion diseases are transmissible and fatal neurodegenerative disorders that affect both human and animals. This disease category comprises Creutzfeldt-Jakob disease (CJD), fatal familial insomnia (FFI) and Gerstmann-Sträussler-Scheinker syndrome (GSS) in humans, scrapie in sheep and goat, bovine spongiform encephalopathy (BSE) in cattle and chronic wasting disease (CWD) in cervids [1]. Prion diseases are thus far incurable. The infectious agent mainly consists of scrapie prion protein (PrP^Sc^), which is a misfolded isoform of the host-encoded cellular prion protein (PrP^C^). PrP^Sc^ acts as a propagon, which can seed a self-perpetuating reaction to recruit and convert PrP^C^ to an aggregated conformation. The deposition of PrP^Sc^ in the central nervous system (CNS), together with neuronal loss, spongiform changes (or termed vacuolization), astrogliosis and conspicuous microglial activation, constitute the characteristic molecular pathology of prion disease [2].

Microglia are the primary innate immune cells and phagocytes of the CNS, exerting a neuroprotective role in prion pathogenesis [3]. We have reported that pharmacogenetic removal of microglia greatly enhances PrP^Sc^ accumulation in prion-infected cultured organotypic cerebellar slices (COCS) and in mice [4, 5]. However, the molecular mechanisms underlying prion clearance by microglia are largely unknown [1]. Lack of milk-fat globule EGF factor VIII (Mfge8) was reported to enhance prion pathogenesis in a mouse strain-dependent manner, suggesting that Mfge8 can facilitate engulfment of PrP^Sc^ aggregates by microglia [6, 7]. Developmental endothelial locuse1 (Del-1) is a structural and functional homolog of Mfge8, and therefore was a further candidate modifier of prion removal. However, Del-1 deficiency neither changed prion deposition nor prion pathogenesis in mice, suggesting that Del-1 does not complement Mfge8 in prion clearance [8]. Also triggering receptor expressed on myeloid cells 2 (TREM2), a phagocytic receptor expressed on microglia, modulates prion-induced microglial activation but does not contribute to prion clearance [9]. Hence, the molecules that are involved in prion clearance are complex and require further study.

Microglia express various receptors that facilitate sensing and phagocytosis of pathogens and misfolded protein aggregates, these include toll-like receptors(TLRs), scavenger receptors(SRs), Fc receptors, complement receptors, triggering receptor expressed on myeloid cells-2 (TREM2), myeloid cell surface antigen CD33 and others [10]. Importantly, variants of the *TREM2* and *CD33* genes are risk factors for Alzheimer’s disease (AD) [11-13], probably due to impaired uptake and clearance of amyloid-β (Aβ) [14, 15]. Although TREM2 is not a main transducer of prion clearance [9], the role of other microglial receptors in prion pathogenesis merits further investigations.

Msr1, also known as scavenger receptor a1 (Scara1), is a type II transmembrane glycoprotein mainly expressed by microglia in CNS [16]. Msr1 has been involved in many macrophage-associated physiological and pathological conditions such as neurodegenerative diseases [17]. As an important phagocytic receptor, Msr1 can mediate uptake of fibrillary amyloid β (Aβ) in vitro [18, 19] and Msr1 deficiency in mouse models of AD markedly accelerates Aβ accumulation and disease progression, whereas pharmacological upregulation of Msr1 leads to enhanced Aβ clearance. These results collectively suggest that Msr1 is essential for clearing soluble Aβ [20, 21]. Since both Aβ and prion are extracellular misfolded protein aggregates, they may share similar molecular pathways by which microglia take up and degrade the protein aggregates.

In this study, we aimed to investigate whether Msr1 may have a similar function in prion clearance and pathogenesis. We first determined the Msr1 expression in a mouse model of prion disease and found Msr1 expression was significantly increased in prion-infected mouse brain. After prion inoculation, we then observed that *Msr1*^-/-^ mice showed disease progression similar to their hemizygous (*Msr1*^+/-^) and wild-type (*Msr1*^+/+^) littermates. Besides, prion deposition and seeding dose were not altered by Msr1 deficiency, suggesting that Msr1 is not involved in prion clearance. Furthermore, Msr1 deficiency did not affect prion-induced neuroinflammation. We therefor conclude that Msr1 is not a major player in prion clearance and does not influence prion pathogenesis.

## Material and methods

### Ethical statement

All animal experiments were carried out in strict accordance with the Rules and Regulations for the Protection of Animal Rights (Tierschutzgesetz and Tierschutzverordnung) of the Swiss Bundesamt für Lebensmittelsicherheit und Veterinärwesen and were preemptively approved by the Animal Welfare Committee of the Canton of Zürich (permit # 41/2012).

### Animals

*Msr1*^-/-^ mice were generated by inserting a neomycin cassette into the EcoRI site in exon 4, which encodes the alpha helical coiled-coil structure essential for the formation of functional trimeric receptors ([22]; JAX stock #006096). *Msr1*^-/-^ mice were first backcrossed to C57BL/6J mice to obtain *Msr1*^+/-^ offspring, which were then intercrossed to generate *Msr1*^+/+^ (wild type), *Msr1*^+/-^ and *Msr1*^-/-^ mice for experiments described here. All animals were maintained in high hygienic grade facility under a 12 h light/12 h dark cycle (from 7 am to 7 pm) at 21±1°C, and fed with diet and water ad libitum.

### Prion inoculation

Mice were intracerebrally (i.c) inoculated with 30 μl of brain homogenate diluted in PBS with 5% BSA and containing 3 × 10^5^ LD50 units of the Rocky Mountain Laboratories scrapie strain (passage 6, thus called RML6). Mice were monitored and actions were taken to minimize animal suffering and distress according to details described previously [23]. Scrapie was diagnosed according to clinical criteria (ataxia, limb weakness, front leg paresis and rolling). Mice were sacrificed by CO2 inhalation on the day of appearance of terminal clinical signs of scrapie (specific criteria referred to [23]), organs were taken and then were either snap-frozen for biochemical analysis or fixed in 4% formalin for histological assessment. The time elapsed from prion inoculation to the terminal stage of disease was defined as incubation time for the survival study.

### Quantitative real-time PCR (qRT-PCR)

Total RNA from was extracted using TRIzol (Invitrogen Life Technologies) according to the manufacturer’s instruction. The quality of RNA was analyzed by Bioanalyzer 2100 (Agilent Technologies), RNAs with RIN>8 were used for cDNA synthesis. cDNAs were synthesized from ∼1 µg total RNA using QuantiTect Reverse Transcription kit (QIAGEN) according to the manufacturer’s instruction. Quantitative real-time PCR (qRT-PCR) was performed using the SYBR Green PCR Master Mix (Roche) on a ViiA7 Real-Time PCR system (Applied Biosystems). The following primer pairs were used: GAPDH sense 5’-CCA CCC CAG CAA GGA GAC T-3’; antisense, 5’-GAA ATT GTG AGG GAG ATG CT-3’. Msr1 sense 5’-TGA ACG AGA GGA TGC TGA CTG -3’; antisense, 5’-GGA GGG GCC ATT TTT AGT GC -3’. TNFα sense, 5’-CAT CTT CTC AAA ATT CGA GTG ACA A-3’; antisense, 5’-TGG GAG TAG ACA AGG TAC AAC CC-3’. IL-1β sense, 5’-CAA CCA ACA AGT GAT ATT CTC CAT G-3’; antisense, 5’-GAT CCA CAC TCT CCA GCT GCA-3’. IL-6 sense, 5’-TCC AAT GCT CTC CTA ACA GAT AAG-3’; antisense, 5’-CAA GAT GAA TTG GAT GGT CTT G -3’. Expression levels were normalized using GAPDH.

### Immunohistochemistry

Prion-infected brain tissues were harvested and fixed in formalin, followed by treatment with concentrated formic acid for 60 min to inactivate prion infectivity and embedded in paraffin. Paraffin sections (2 μm) of brains were stained with hematoxylin/eosin (HE) to visualize prion-induced lesions and vacuolation. For the histological detection of partially proteinase K-resistant prion protein deposition, deparaffinized sections were pretreated with formaldehyde for 30 min and 98% formic acid for 6 min, and then washed in distilled water for 30 min. Sections were incubated in Ventana buffer and stains were performed on a NEXEX immunohistochemistry robot (Ventana instruments, Switzerland) using an IVIEW DAB Detection Kit (Ventana). After incubation with protease 1 (Ventana) for 16 min, sections were incubated with anti-PrP SAF-84 (1:200, SPI bio, A03208) for 32 min. Sections were counterstained with hematoxylin. To detect astrogliosis and microglial activation, brain sections were deparaffinized through graded alcohols, anti-GFAP antibody (1:300; DAKO, Carpinteria, CA) were applied for astrogliosis, anti-Iba1 antibody (1:1000; Wako Chemicals GmbH, Germany) was used for highlighting activated microglial cells. Stainings were visualized using DAB (Sigma-Aldrich), hematoxylin counterstain was subsequently applied. Sections were imaged using a Zeiss Axiophot light microscope.

### Western blot analysis

To detect PrP^C^ in the mouse brains, one hemisphere of each brain was homogenized with RIPA buffer. Total protein concentration was determined using the bicinchoninic acid assay (Pierce). ∼8 ug proteins were loaded and separated on a 12% Bis-Tris polyacrylamide gel (NuPAGE, Invitrogen) and then blotted onto a nitrocellulose membrane. Membranes were blocked with 5% wt/vol Topblock (LuBioScience) in PBS supplemented with 0.1% Tween 20 (vol/vol) and incubated with primary antibodies POM1 in 1% Topblock (400 ng ml^−1^) overnight. After washing, the membranes were then incubated with secondary antibody horseradish peroxidase (HRP)-conjugated goat anti–mouse IgG (1:10,000, Jackson ImmunoResearch, 115-035-003). Blots were developed using Luminata Crescendo Western HRP substrate (Millipore) and visualized using the Stella system (model 3000, Bio-Rad). To avoid variation in loading, the same blots were stripped and incubated with anti-actin antibody (1:10,000, Millipore). The PrP^C^ signals were normalized to actin as a loading control.

To detect PrP^Sc^, prion infected forebrains were homogenized in sterile 0.32 M sucrose in PBS. Total protein concentration was determined using the bicinchoninic acid assay (Pierce). Samples were adjusted to 20 µg protein in 20 µl and digested with 25 µg ml^−1^ proteinase K for 30 min at 37°C. PK digestion was stopped by adding loading buffer (Invitrogen) and boiling samples at 95°C for 5 min. Proteins were then separated on a 12% Bis-Tris polyacrylamide gel (NuPAGE, Invitrogen) and blotted onto a nitrocellulose membrane. POM1 and horseradish peroxidase (HRP)-conjugated goat anti–mouse IgG were used as primary and secondary antibodies, respectively. Blots were developed using Luminata Crescendo Western HRP substrate (Millipore) and visualized using the FUJIFILM LAS-3000 system. To detect GFAP and Iba1 in prion-infected brains by Western blot, 20 µg of total brain protein were loaded and anti-GFAP antibody (D1F4Q) XP Rabbit mAb (1:3000; Cell Signaling Technology, 12389), anti-Iba1 antibody (1:1000; Wako Chemicals GmbH, Germany, 019-19741) and horseradish peroxidase (HRP)-conjugated goat anti-rabbit IgG (1:10,000, Jackson ImmunoResearch, 111-035-045) were used as primary and secondary antibodies, respectively. Actin was used as the loading control.

### Real-time quaking induced conversion assay (RT-QuIC)

RT-QuIC assays of prion-infected mouse brain homogenates were performed as previously described [8, 24]. Briefly, recombinant hamster full-length (23–231) PrP was expressed in Rosetta2(DE3)pLysS E.coli competent cells and purified by affinity chromatography using Ni2^+^-nitrilotriacetic acid Superflow resin (QIAGEN). In the RT-QuIC assay, recombinant HaPrP was used as substrate for PrP^Sc^-catalyzed conversion. RT-QuIC reactions containing HaPrP substrate protein at a final concentration of 0.1 mg mL^-1^ in PBS (pH 7.4), 170 mM NaCl, 10 μM EDTA, 10 μM Thioflavin T were seeded with 2 μL of serially diluted brain homogenates in a total reaction volume of 100 μL. NBH- and RML6-treated brain homogenates were used as negative and positive controls, respectively. The RT-QuIC reactions were amplified at 42 °C for 100 h with intermittent shaking cycles of 90 s shaking at 900 rpm in double orbital mode and 30 s resting using a FLUOstar Omega microplate reader (BMG Labtech). Aggregate formation was followed by measuring the thioflavin T fluorescence every 15 min (450 nm excitation, 480 nm emission; bottom read mode).

### Statistical analysis

Results are presented as the mean of replicas ± standard error of the mean (SEM). Statistical significance between experimental groups was assessed using an unpaired Student’s t-Test. For prion inoculation experiments, incubation times were analyzed using the Kaplan-Meier method and compared between groups using Log-rank (Mantel-Cox) test. p-values <0.05 were considered statistically significant.

## Results

### Prion infection upregulates Msr1 expression in the mouse brain

Expression of Msr1 could be upregulated by various forms of brain injury [25, 26]. To test whether prion infection affects Msr1 expression in the mouse brain, we intracerebrally (i.c) inoculated 30 µl of diluted CD1 mouse brain homogenate containing 3 × 10^5^ LD50 (50% lethal dose) units of RML6 (a prion strain originated from the Rocky Mountain Laboratory, serially passaged to No. 6, hence termed RML6) into C57BL/6J mice. C57BL/6J littermates inoculated with noninfectious brain homogenate (NBH) were used as control. Prion-inoculated mice were euthanized and brains were collected when they showed severe scrapie sign and reached the terminal stage of disease, NBH-inoculated control mice were sacrificed after the same incubation time. Since we could not commercially obtain a sensitive and specific antibody detecting the Msr1 protein in mouse brains (supplemental figure 1), we assessed Msr1 expression levels using quantitative reverse-transcription PCR (qRT-PCR). The results showed that Msr1 expression was significantly enhanced in prion-inoculated mouse brain, compared to NBH inoculated mouse (Figure 1A), suggesting that Msr1 expression can be efficiently upregulated by prion infection.

**Figure 1:**
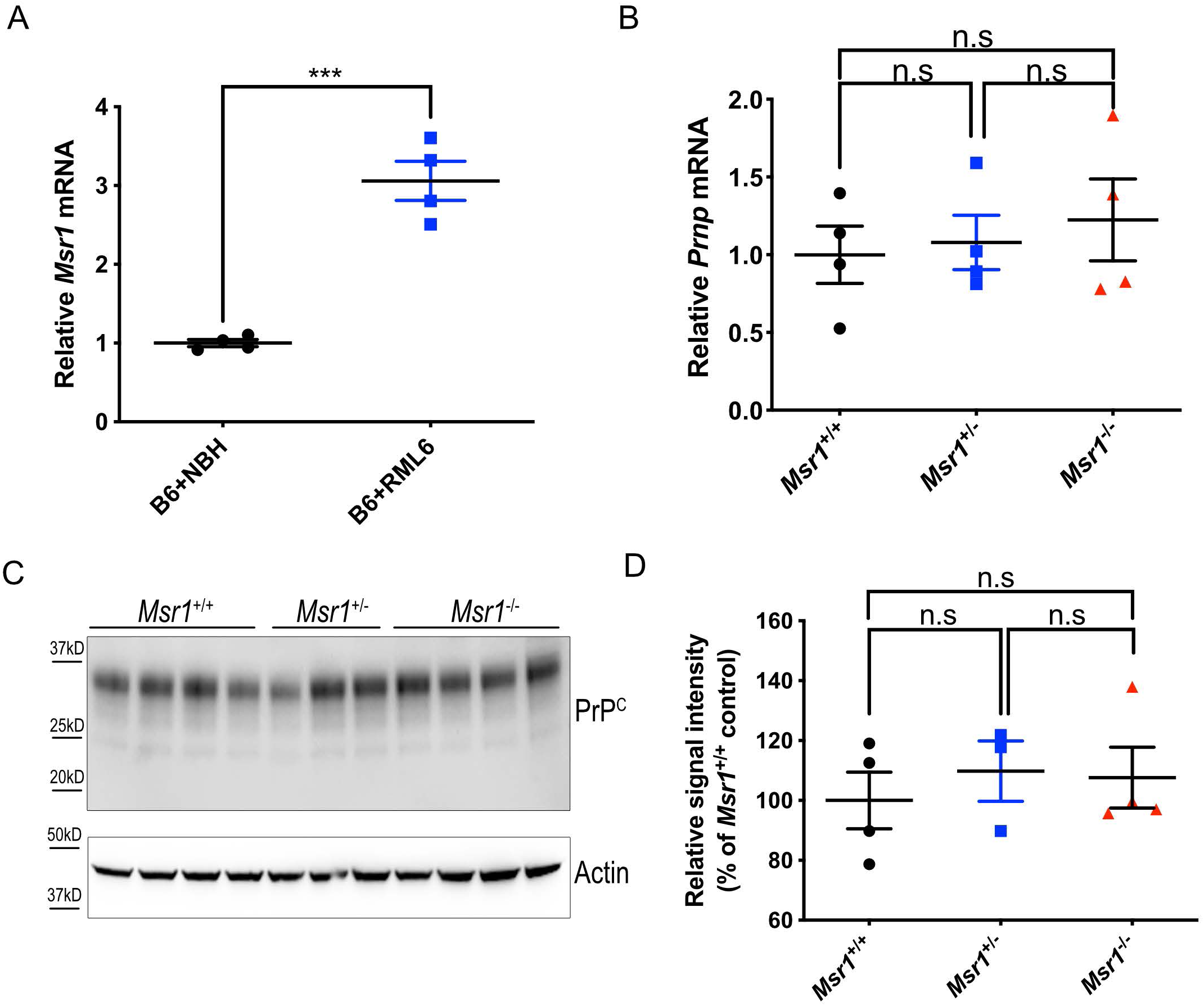
(A) qRT-PCR of Msr1 expression in mouse brains. Msr1 expression was significantly increased in RML inoculated mouse brain, compared to NBH inoculated mouse brain (n=4, p<0.001). (B) *Prnp* qRT-PCR in *Msr1*^+/+^ (WT), *Msr1*^+/-^ and *Msr1*^-/-^ mouse brains (n=4, n.s p>0.05). No significant difference of *Prnp* mRNA was observed between three groups. (C and D) PrP^C^ Western blot (C) and densitometric quantification of the Western blot (D) in *Msr1*^+/+^ (WT), *Msr1*^+/-^ and *Msr1*^-/-^ mouse brains (n=3∼4, n.s p>0.05). No significant difference of PrP^C^ protein was observed between three groups. Abbreviations: Msr1, macrophage scavenger receptor 1; *Prnp*, murine prion gene; PrP^C^, cellular prion protein; qRT-PCR, quantitative real-time polymerase chain reaction; RML6, Rocky Mountain Laboratories scrapie strain, passage 6; NBH, noninfectious brain homogenates; WT, wild type.

PrP^C^ expression levels in mouse brains are the major determinants of the susceptibility to prion diseases and progression rate of the diseases [27, 28]. To assess whether Msr1 deficiency could influence PrP^C^ expression, we performed qRT-PCR and Western blot to measure *Prnp* mRNA transcripts and PrP^C^ protein levels in *Msr1*^+/+^, *Msr1*^+/-^ and *Msr1*^-/-^ littermates. The results showed that Msr1 deficiency altered neither the *Prnp* mRNA (Figure 1B) nor the protein levels of PrP^C^ (Figure 1C-D, uncropped Western blots are shown in Supplementary Figure 2A-2B) in mouse brains.

### Prion disease progression, lesion pattern or PrP^Sc^ accumulation are not altered by Msr1 deficiency

To assess the function of Msr1 in prion pathogenesis, we next tested whether Msr1 ablation could affect prion disease progression and alter prion-mediated lesion pattern in mouse brains. We intracerebrally (i.c) inoculated 30 µl of RML6 prions into *Msr1*^+/+^, *Msr1*^+/-^ and *Msr1*^-/-^ littermates. RML6-inoculated mice were checked and monitored every other day for scrapie symptoms. Mice were euthanized and brains were collected when they showed severe scrapie sign and reached the terminal stage of disease. Incubation times were calculated as the time from initial prion inoculation until terminal disease stage. We observed that all *Msr1*^+/+^, *Msr1*^+/-^ and *Msr1*^-/-^ mice succumbed to prion disease at a similar progression rate (median survival: 179 dpi for *Msr1*^+/+^ mice (n=17), 186 dpi for *Msr1*^+/-^(n=25) and 183.5 dpi for *Msr1*^-/-^ mice (n=14), p=0.99) (Figure 2A). The above results indicate that Msr1 ablation does not overtly influence progression of prion disease.

**Figure 2:**
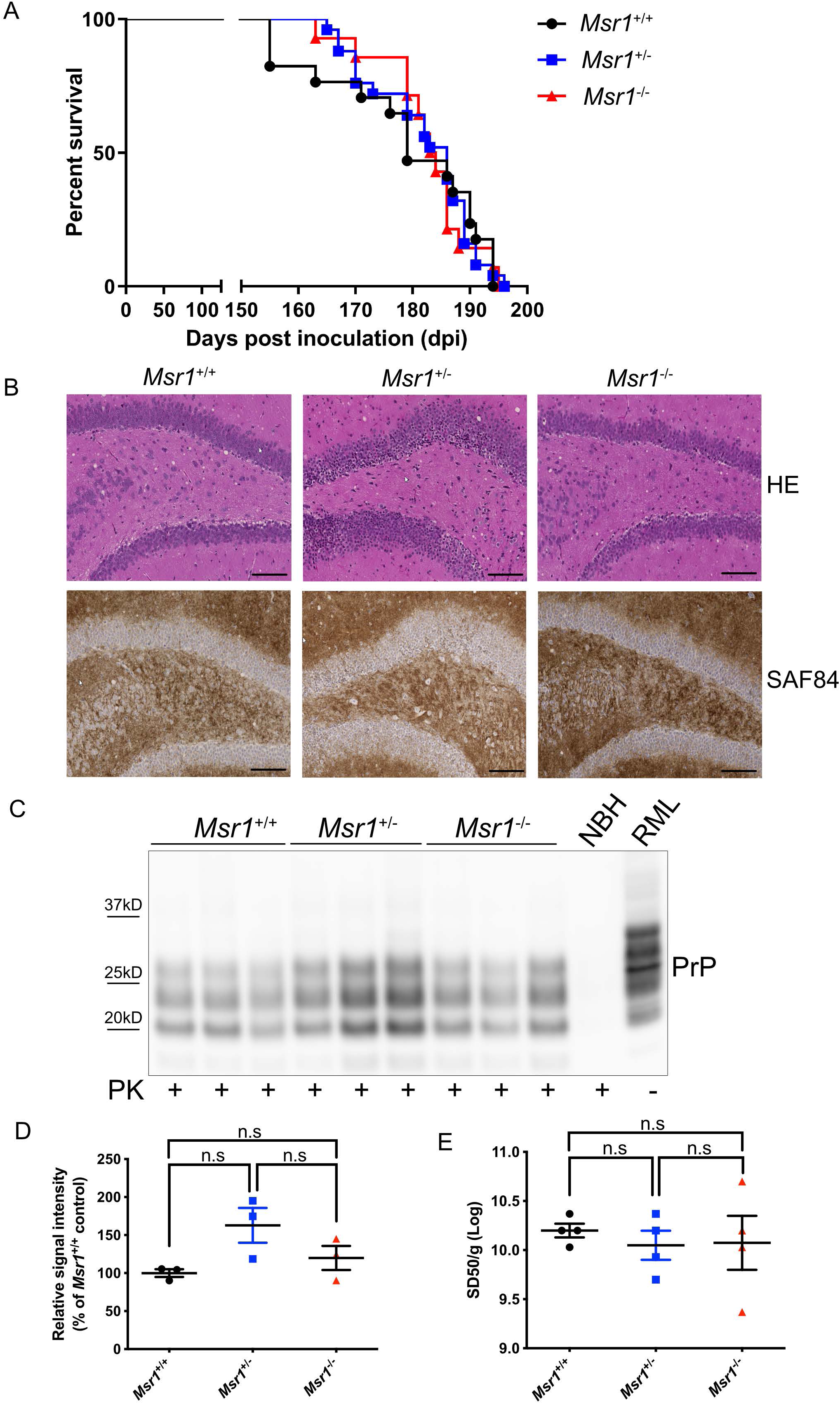
(A) Survival curve of *Msr1*^+/+^ (WT), *Msr1*^+/-^ and *Msr1*^-/-^ littermates intracerebrally inoculated with RML6. There was no significant difference between three groups in both genders (n=14∼25, n.s *p*>0.05). (B) Representative histology of terminally sick mouse brains from *Msr1*^+/+^ (WT), *Msr1*^+/-^ and *Msr1*^-/-^ littermates stained for H&E and SAF84. There was no obvious difference between three groups in vacuolation, lesion pattern and PrP^Sc^ deposition in hippocampus. (C and D) Western blot for proteinase K resistant PrP^Sc^ in terminally sick mouse brains (C) and densitometric quantification of the Western blot (D). There was no significantly difference between *Msr1*^+/+^ (WT), *Msr1*^+/-^ and *Msr1*^-/-^ littermates (n=3, n.s *p*>0.05). (E) RT-QuIC assay of terminally sick mouse brains from *Msr1*^+/+^ (WT), *Msr1*^+/-^ and *Msr1*^-/-^ littermates showed similar level of 50% prion seeding dose in these three groups (n=4, n.s *p*>0.05). Abbreviations: Msr1, macrophage scavenger receptor 1; PrP^Sc^, scrapie-associated prion protein; RML6, Rocky Mountain Laboratories scrapie strain, passage 6; NBH, noninfectious brain homogenates; WT, wild type; RT-QuIC, real-time quaking-induced conversion assay.

We then analyzed and compared the histology of brain sections collected and prepared from RML6-inoculated terminally sick *Msr1*^+/+^, *Msr1*^+/-^ and *Msr1*^-/-^ mice. The classical histological characteristics of prion disease, especially the spongiform changes (or vacuolation), were observed in all mice in different groups. Lesion pattern analysis also failed to show any qualitative distinctions between different genotypes (Figure 2B). These results suggest that lack of Msr1 does not obviously affect prion-caused lesion profile in mouse brains.

If Msr1 contributed to prion clearance as it does to Aβ, *Msr1*^-/-^ mice would accumulate more PrP^Sc^ deposits in their brains. We therefore first performed PrP^Sc^ staining on brain sections prepared from RML6-inoculated terminally sick *Msr1*^+/+^, *Msr1*^+/-^ and *Msr1*^-/-^ mice. Unexpectedly, we observed a similar PrP^Sc^ deposition level and pattern in mouse brains with the different genotypes (Figure 2B). We next performed Western blot to detect and assess proteinase K (PK)-resistant PrP^Sc^ levels in brains of terminally sick mice. We again found a similar PrP^Sc^ accumulation level in *Msr1*^+/+^, *Msr1*^+/-^ and *Msr1*^-/-^ mouse brains (Figure 2C-D, uncropped Western blots are shown in Supplementary Figure 2C). Additionally, we measured the prion seeding activity in brain homogenates of RML6-inoculated terminally sick *Msr1*^+/+^, *Msr1*^+/-^ and *Msr1*^-/-^ mice by Real Time-Quaking Induced Conversion assay (RT-QuIC). We detected an undistinguishable 50% seeding dose (SD50) of prion in all three groups (Figure 2E). Taken together, these results indicate that Msr1 does not play a major role in prion clearance or PrP^Sc^ accumulation in mouse brains.

### Prion-induced neuroinflammation is not affected by Msr1 deficiency

Msr1 is a scavenger receptor that could modulate immune response in CNS [29-31]. To test whether Msr1 exhibits an anti-inflammatory function in prion pathogenesis, we analyzed and compared astrogliosis and microglial activation in prion-infected terminally sick *Msr1*^+/+^, *Msr1*^+/-^ and *Msr1*^-/-^ mice. First, histology and Western blots did not identify obvious differences in GFAP (glial fibrillary acidic protein, an astrocytic marker) level between the three groups with different genotypes (Figure 3A-B, uncropped Western blots are shown in Supplementary Figure 2D-2E). We next performed histological examinations and Western blots to detect Iba1 (ionized calcium binding adaptor molecule 1, a microglial marker) in RML6-inoculated terminally sick *Msr1*^+/+^, *Msr1*^+/-^ and *Msr1*^-/-^ mouse brains. We again observed similar levels of Iba1 between the three groups (Figure 3C-D, uncropped Western blots are shown in Supplementary Figure 2F-G). These results together suggest that Msr1 deficiency does not significantly affect prion-induced astrogliosis or microglial activation.

**Figure 3:**
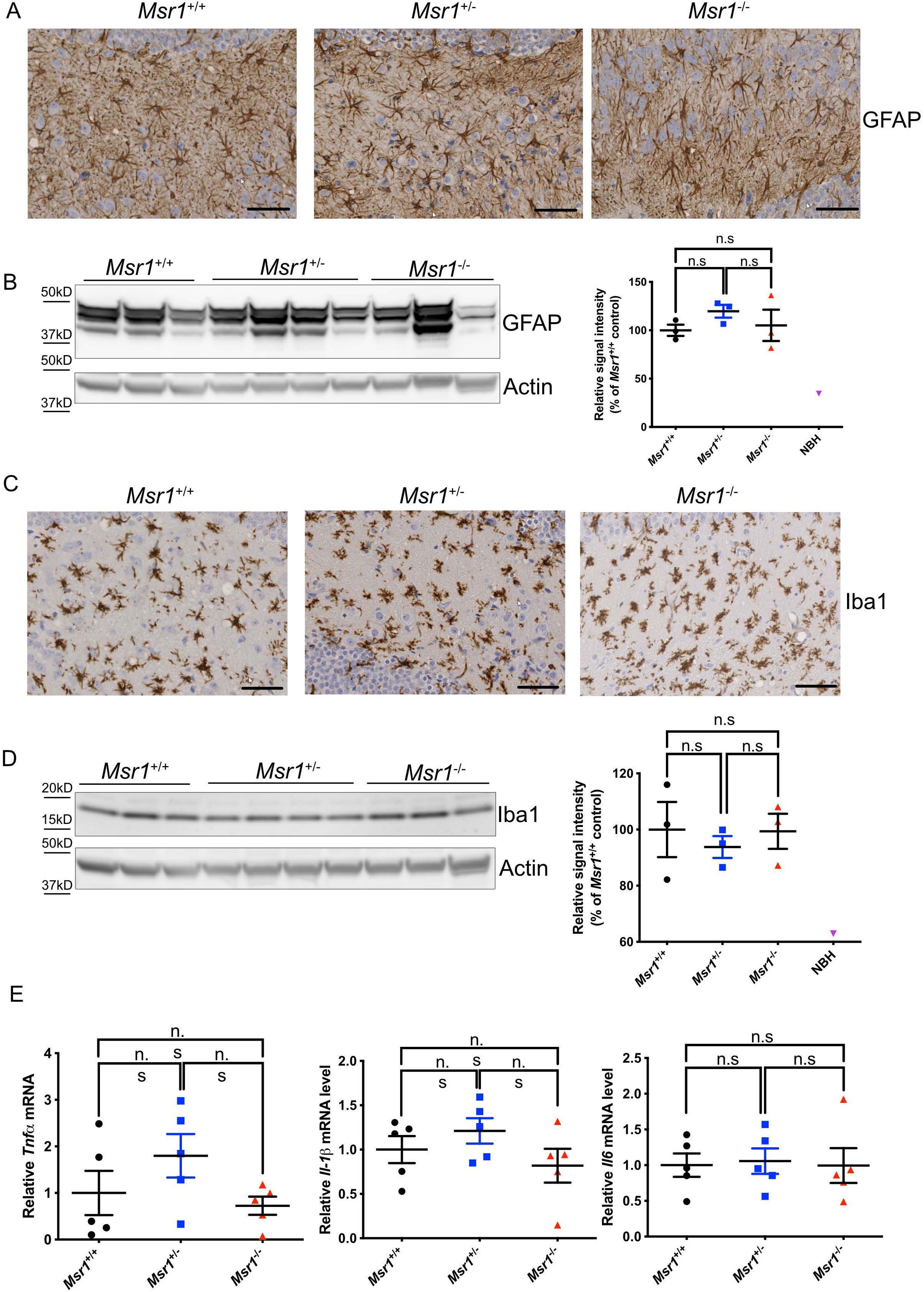
(A) Representative immunohistochemical staining for GFAP in the hippocampus of terminally sick mouse brains from *Msr1*^+/+^ (WT), *Msr1*^+/-^ and *Msr1*^-/-^ littermates. (B) Left: Western blot for GFAP in terminally sick mouse brains. Right: densitometric quantification of the Western blot showed no significant difference of GFAP levels between *Msr1*^+/+^ (WT), *Msr1*^+/-^ and *Msr1*^-/-^ littermates (n=3∼4, n.s *p*>0.05). (C) Representative immunohistochemical staining for Iba1 in the hippocampus of terminally sick mouse brains from *Msr1*^+/+^ (WT), *Msr1*^+/-^ and *Msr1*^-/-^ littermates. (D) Left: Western blot for Iba1 in terminally sick mouse brains. Right: densitometric quantification of the Western blot showed no significant difference of Iba1 levels between *Msr1*^+/+^ (WT), *Msr1*^+/-^ and *Msr1*^-/-^ littermates (n=3∼4, n.s *p*>0.05). (E) qRT-PCR of cytokines TNFα, IL-1β and IL-6 expression revealed similar mRNA levels of these cytokines (*Tnfα, Il1β,Il6*) in terminally sick *Msr1*^+/+^ (WT), *Msr1*^+/-^ and *Msr1*^-/-^ littermates (n=5, n.s *p*>0.05). Abbreviations: Msr1, macrophage scavenger receptor 1; GFAP, glial fibrillary acidic protein; Iba1, Ionized calcium binding adaptor molecule 1; WT, wild type; qRT-PCR, quantitative real-time polymerase chain reaction.

We next performed qRT-PCR to analyze the expression levels of proinflammatory cytokines (TNFα, IL-1β and IL6) in RML6-inoculated terminally sick *Msr1*^+/+^, *Msr1*^+/-^ and *Msr1*^-/-^ mouse brains. Again, this experiment failed to detect any overt differences in cytokine expression levels between the three groups (Figure 3E), suggesting that Msr1 ablation does not change the cytokine profiles of prion-infected mouse brains.

## Discussion

It is becoming increasingly clear that neuroinflammation, characterized by astrogliosis and microglial activation, is a hallmark of many neurodegenerative diseases including prion diseases [32]. Emerging evidence from multiple genome-wide association studies (GWAS) suggests that microglia may play a central role in neurodegenerative diseases because most identified risk factors are expressed by microglia [11, 12, 33-37]. The molecular mechanisms by which microglia contributes to the pathogenesis of neurodegenerative diseases are the subject of intense investigations. Insight into these questions will enable not only the understanding of disease pathogenesis, but also the development of novel therapeutic strategies combating these disorders [38].

As the primary tissue-resident macrophages in the CNS, microglia exert diverse functions in the brain. Microglia play important roles during brain development by pruning neuronal synapses [39]. In adult brain, microglia could modulate learning and memory [40], and constantly survey their local milieu for signals of danger and injury, thus serving as important sensors and defenders upon challenges [41, 42]. Under pathological conditions such as in prion disease, microglia are found to play an overall neuroprotective role by clearing prions. Depletion or deficiency of microglia results in impaired prion clearance, enhanced PrP^Sc^ deposition and deteriorated prion pathogenesis [5]. These findings collectively highlight the need to better understand the molecular mechanisms underlying microglia-mediated prion clearance [3].

Msr1, a type II transmembrane glycoprotein expressed mainly by microglia in the brain [16], is a member of scavenger receptors that mediate endocytosis of a wide range of ligands including low-density lipoproteins, bacterial pathogens and dead cells [43, 44]. The role of Msr1 in neurodegeneration such as Alzheimer’s disease (AD) was first highlighted in an *in vitro* study showing that it could facilitate adhesion of microglia to fibrillar Aβ [18]. This finding was further validated by a short hairpin RNA (shRNA) library screen [20]. In mouse models of AD, Msr1 deficiency impairs clearance of soluble Aβ and results in increased Aβ deposition and early mortality, whereas pharmacological upregulation of Msr1 leads to enhanced Aβ clearance [20, 21]. Therefore, it is reasonable to speculate that Msr1 could play a scavenger function for a broad range of misfolded protein aggregates including prions.

In this study, we tested whether Msr1 plays a scavenger function in prion pathogenesis, similar to that in mouse models of AD. Firstly, in contrast to what was observed in aged AD mouse models [45, 46], Msr1 expression was found to be upregulated upon prion infection. After prion inoculation, we observed that *Msr1*^-/-^ mice experienced a similar incubation time compared with *Msr1*^+/-^ and wild type littermates, suggesting that Msr1 deficiency does not affect prion disease progression. Notably, the PrP^Sc^ level and the prion seeding activity were not detectably affected by Msr1 deficiency, indicating that Msr1 is not a major contributor to prion clearance. Moreover, prion-induced neuroinflammation including astrocytosis and microglial activation was not altered in Msr1^-/-^ mouse brains, suggesting that Msr1 deficiency does not overtly affect prion-induced neuroinflammation. Together, these results indicate that Msr1 functions differently in AD and prion diseases.

Collectively, these results indicate that Msr1 does not play a major role in prion pathogenesis. Together with the discrepant observations of TREM2 in AD and prion disease[9, 14], this study suggests that microglia-mediated phagocytosis and clearance of Aβ and prion may adopt distinct molecular pathways. Further studies are needed to investigate molecular mechanisms underlying microglial uptake and clearance of prions.

## Acknowledgements

We thank the teams of the Institute of Neuropathology, University Hospital Zurich and Department of Neurobiology, School of Basic Medical Sciences of Fudan University for technical assistance. The authors thank Elisabeth J. Rushing for reading and editing the article. A. Aguzzi is the recipient of an Advanced Grant of the European Research Council (ERC, No. 670958) and is supported by grants from the European Union (PRIORITY, NEURINOX), the Swiss National Foundation (SNF, including a Sinergia grant), the Swiss Initiative in Systems Biology, SystemsX.ch (PrionX, SynucleiX), and a Distinguished Scientist Award of the Nomis Foundation. C. Zhu is sponsored by Research Startup Funds of Fudan University, Shanghai Pujiang Program (No. 20PJ1401100) and National Natural Science Foundation of China (No. 82071436). The funders had no role in study design, data collection and analysis, decision to publish, or preparation of the manuscript.

## Declarations

### Consent to participate

Not applicable.

### Consent to publication

The authors confirm that the manuscript has been read and approved by all named authors. All authors consent to the publication of the manuscript on Journal of Molecular Medicine.

### Availability of data and material

The authors confirm that the data and material supporting the findings of this study are available within the article and its supplementary materials.

### Code availability

Not applicable.

### Authors’ contributions

C. Zhu and A. Aguzzi conceived the project and designed the experiments; B. Li, M. Chen and C. Zhu conducted the experiments and acquired data. All authors contributed to data analysis.

---

**Supplemental figure 1:**
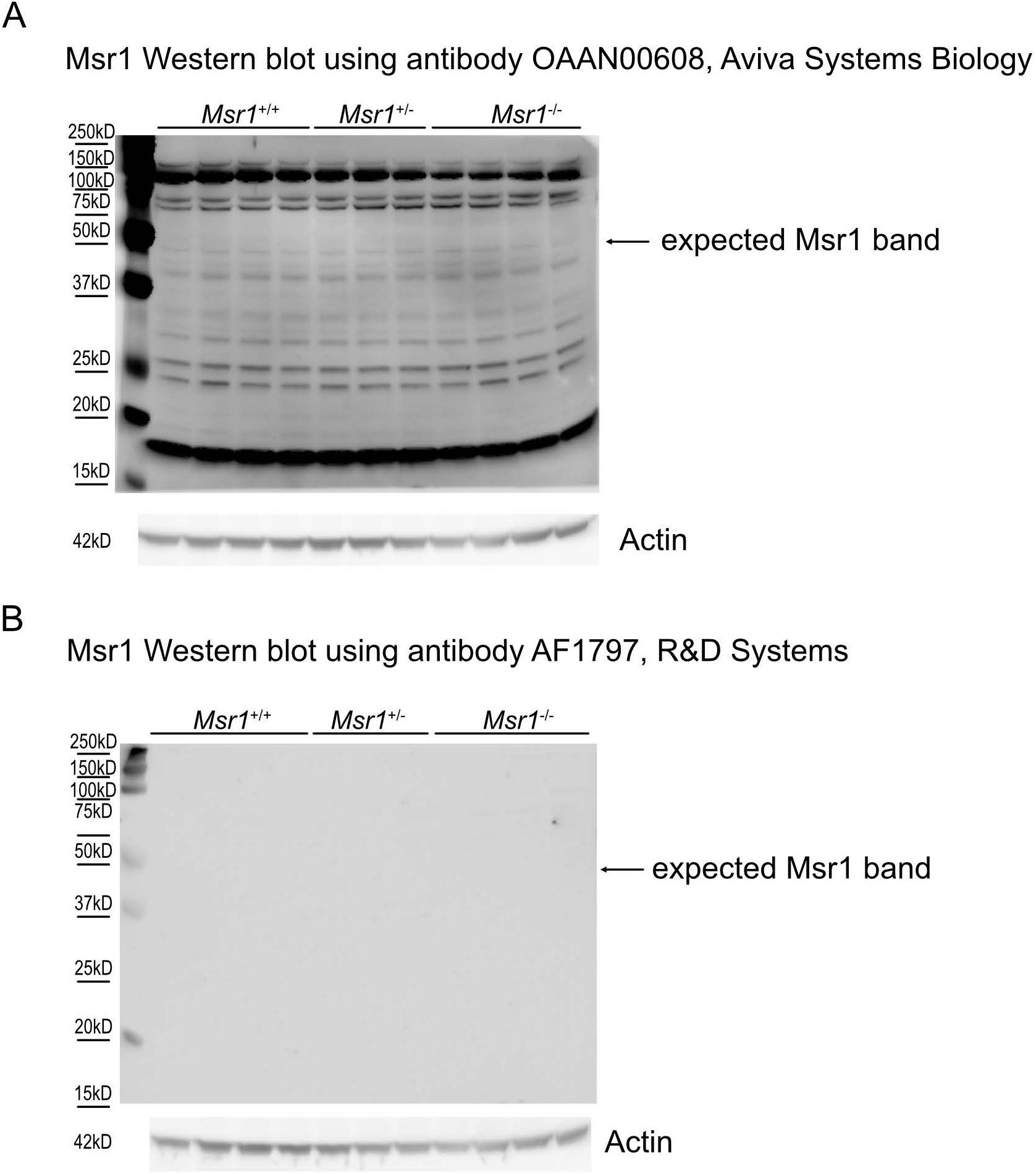
Western blots using commercial anti-Msr1 antibodies that could not detect specific Msr1 band on brains collected from wild type mice. *Msr1*^-/-^ mouse brains were used as negative control.

**Supplemental figure 2:**
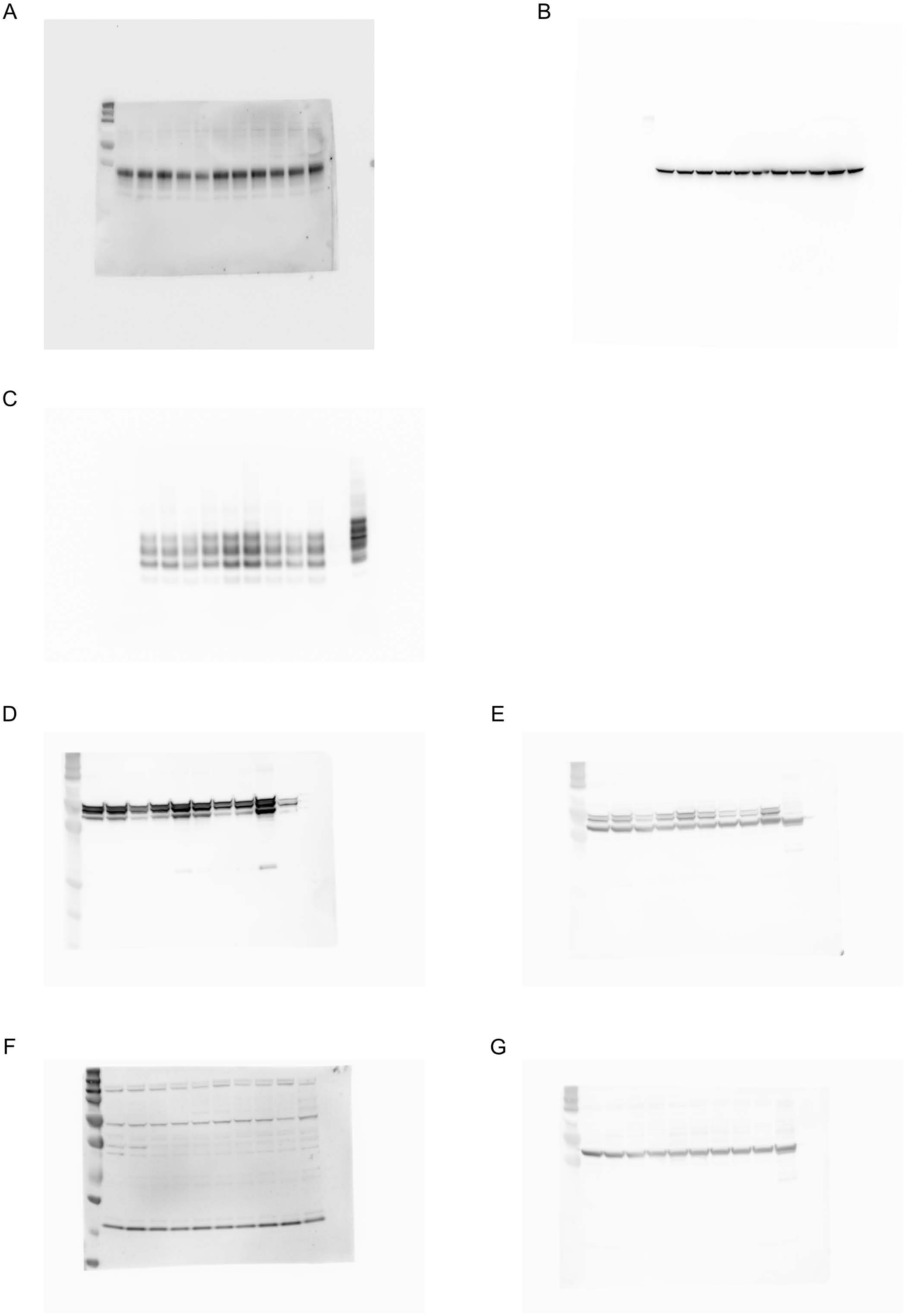
Full images of the cropped Western blots in figure 1D (A-B), 2C (C), 3B (D-E) and 3D (F-G).

## References

1. Aguzzi A, Nuvolone M, Zhu C (2013) The immunobiology of prion diseases. Nat Rev Immunol 13: 888–902. DOI 10.1038/nri3553

2. Aguzzi A, Zhu C (2012) Five questions on prion diseases. PLoS Pathog 8: e1002651. DOI 10.1371/journal.ppat.1002651

3. Aguzzi A, Zhu C (2017) Microglia in prion diseases. J Clin Invest 127: 3230–3239. DOI 10.1172/JCI90605

4. Falsig J, Julius C, Margalith I, Schwarz P, Heppner FL, Aguzzi A (2008) A versatile prion replication assay in organotypic brain slices. Nat Neurosci 11: 109–117. DOI 10.1038/nn2028

5. Zhu C, Herrmann US, Falsig J, Abakumova I, Nuvolone M, Schwarz P, Frauenknecht K, Rushing EJ, Aguzzi A (2016) A neuroprotective role for microglia in prion diseases. J Exp Med 213: 1047–1059. DOI 10.1084/jem.20151000

6. Kranich J, Krautler NJ, Heinen E, Polymenidou M, Bridel C, Schildknecht A, Huber C, Kosco-Vilbois MH, Zinkernagel R, Miele G, et al. (2008) Follicular dendritic cells control engulfment of apoptotic bodies by secreting Mfge8. J Exp Med 205: 1293–1302. DOI 10.1084/jem.20071019

7. Kranich J, Krautler NJ, Falsig J, Ballmer B, Li S, Hutter G, Schwarz P, Moos R, Julius C, Miele G, et al. (2010) Engulfment of cerebral apoptotic bodies controls the course of prion disease in a mouse strain-dependent manner. J Exp Med 207: 2271–2281. DOI 10.1084/jem.20092401

8. Zhu C, Li Z, Li B, Pfammatter M, Hornemann S, Aguzzi A (2019) Unaltered prion disease in mice lacking developmental endothelial locus-1. Neurobiol Aging 76: 208–213. DOI 10.1016/j.neurobiolaging.2019.01.003

9. Zhu C, Herrmann US, Li B, Abakumova I, Moos R, Schwarz P, Rushing EJ, Colonna M, Aguzzi A (2015) Triggering receptor expressed on myeloid cells-2 is involved in prion-induced microglial activation but does not contribute to prion pathogenesis in mouse brains. Neurobiol Aging 36: 1994–2003. DOI 10.1016/j.neurobiolaging.2015.02.019

10. Doens D, Fernandez PL (2014) Microglia receptors and their implications in the response to amyloid beta for Alzheimer’s disease pathogenesis. J Neuroinflammation 11: 48. DOI 10.1186/1742-2094-11-48

11. Guerreiro R, Wojtas A, Bras J, Carrasquillo M, Rogaeva E, Majounie E, Cruchaga C, Sassi C, Kauwe JS, Younkin S, et al. (2013) TREM2 variants in Alzheimer’s disease. N Engl J Med 368: 117–127. DOI 10.1056/NEJMoa1211851

12. Jonsson T, Stefansson H, Steinberg S, Jonsdottir I, Jonsson PV, Snaedal J, Bjornsson S, Huttenlocher J, Levey AI, Lah JJ, et al. (2013) Variant of TREM2 associated with the risk of Alzheimer’s disease. N Engl J Med 368: 107–116. DOI 10.1056/NEJMoa1211103

13. Bradshaw EM, Chibnik LB, Keenan BT, Ottoboni L, Raj T, Tang A, Rosenkrantz LL, Imboywa S, Lee M, Von Korff A, et al. (2013) CD33 Alzheimer’s disease locus: altered monocyte function and amyloid biology. Nat Neurosci 16: 848–850. DOI 10.1038/nn.3435

14. Wang Y, Cella M, Mallinson K, Ulrich JD, Young KL, Robinette ML, Gilfillan S, Krishnan GM, Sudhakar S, Zinselmeyer BH, et al. (2015) TREM2 lipid sensing sustains the microglial response in an Alzheimer’s disease model. Cell 160: 1061–1071. DOI 10.1016/j.cell.2015.01.049

15. Griciuc A, Serrano-Pozo A, Parrado AR, Lesinski AN, Asselin CN, Mullin K, Hooli B, Choi SH, Hyman BT, Tanzi RE (2013) Alzheimer’s disease risk gene CD33 inhibits microglial uptake of amyloid beta. Neuron 78: 631–643. DOI 10.1016/j.neuron.2013.04.014

16. Christie RH, Freeman M, Hyman BT (1996) Expression of the macrophage scavenger receptor, a multifunctional lipoprotein receptor, in microglia associated with senile plaques in Alzheimer’s disease. Am J Pathol 148: 399–403

17. Wilkinson K, El Khoury J (2012) Microglial scavenger receptors and their roles in the pathogenesis of Alzheimer’s disease. Int J Alzheimers Dis 2012: 489456. DOI 10.1155/2012/489456

18. El Khoury J, Hickman SE, Thomas CA, Cao L, Silverstein SC, Loike JD (1996) Scavenger receptor-mediated adhesion of microglia to beta-amyloid fibrils. Nature 382: 716–719. DOI 10.1038/382716a0

19. Chung H, Brazil MI, Irizarry MC, Hyman BT, Maxfield FR (2001) Uptake of fibrillar beta-amyloid by microglia isolated from MSR-A (type I and type II) knockout mice. Neuroreport 12: 1151–1154. DOI 10.1097/00001756-200105080-00020

20. Frenkel D, Wilkinson K, Zhao L, Hickman SE, Means TK, Puckett L, Farfara D, Kingery ND, Weiner HL, El Khoury J (2013) Scara1 deficiency impairs clearance of soluble amyloid-beta by mononuclear phagocytes and accelerates Alzheimer’s-like disease progression. Nat Commun 4: 2030. DOI 10.1038/ncomms3030

21. Sandoval K, Umbaugh D, House A, Crider A, Witt K (2019) Somatostatin Receptor Subtype-4 Regulates mRNA Expression of Amyloid-Beta Degrading Enzymes and Microglia Mediators of Phagocytosis in Brains of 3xTg-AD Mice. Neurochem Res 44: 2670–2680. DOI 10.1007/s11064-019-02890-6

22. Suzuki H, Kurihara Y, Takeya M, Kamada N, Kataoka M, Jishage K, Ueda O, Sakaguchi H, Higashi T, Suzuki T, et al. (1997) A role for macrophage scavenger receptors in atherosclerosis and susceptibility to infection. Nature 386: 292–296. DOI 10.1038/386292a0

23. Zhu C, Schwarz P, Abakumova I, Aguzzi A (2015) Unaltered Prion Pathogenesis in a Mouse Model of High-Fat Diet-Induced Insulin Resistance. PLoS One 10: e0144983. DOI 10.1371/journal.pone.0144983

24. Frontzek K, Pfammatter M, Sorce S, Senatore A, Schwarz P, Moos R, Frauenknecht K, Hornemann S, Aguzzi A (2016) Neurotoxic Antibodies against the Prion Protein Do Not Trigger Prion Replication. PLoS One 11: e0163601. DOI 10.1371/journal.pone.0163601

25. Bell MD, Lopez-Gonzalez R, Lawson L, Hughes D, Fraser I, Gordon S, Perry VH (1994) Upregulation of the macrophage scavenger receptor in response to different forms of injury in the CNS. J Neurocytol 23: 605–613. DOI 10.1007/BF01191555

26. Grewal RP, Yoshida T, Finch CE, Morgan TE (1997) Scavenger receptor mRNAs in rat brain microglia are induced by kainic acid lesioning and by cytokines. Neuroreport 8: 1077–1081. DOI 10.1097/00001756-199703240-00003

27. Bueler H, Raeber A, Sailer A, Fischer M, Aguzzi A, Weissmann C (1994) High prion and PrPSc levels but delayed onset of disease in scrapie-inoculated mice heterozygous for a disrupted PrP gene. Mol Med 1: 19–30

28. Fischer M, Rulicke T, Raeber A, Sailer A, Moser M, Oesch B, Brandner S, Aguzzi A, Weissmann C (1996) Prion protein (PrP) with amino-proximal deletions restoring susceptibility of PrP knockout mice to scrapie. EMBO J 15: 1255–1264

29. Peiser L, Mukhopadhyay S, Gordon S (2002) Scavenger receptors in innate immunity. Curr Opin Immunol 14: 123–128. DOI 10.1016/s0952-7915(01)00307-7

30. Areschoug T, Gordon S (2009) Scavenger receptors: role in innate immunity and microbial pathogenesis. Cell Microbiol 11: 1160–1169. DOI 10.1111/j.1462-5822.2009.01326.x

31. Canton J, Neculai D, Grinstein S (2013) Scavenger receptors in homeostasis and immunity. Nat Rev Immunol 13: 621–634. DOI 10.1038/nri3515

32. Aguzzi A, Barres BA, Bennett ML (2013) Microglia: scapegoat, saboteur, or something else? Science 339: 156–161. DOI 10.1126/science.1227901

33. Baker M, Mackenzie IR, Pickering-Brown SM, Gass J, Rademakers R, Lindholm C, Snowden J, Adamson J, Sadovnick AD, Rollinson S, et al. (2006) Mutations in progranulin cause tau-negative frontotemporal dementia linked to chromosome 17. Nature 442: 916–919. DOI 10.1038/nature05016

34. Cruts M, Gijselinck I, van der Zee J, Engelborghs S, Wils H, Pirici D, Rademakers R, Vandenberghe R, Dermaut B, Martin JJ, et al. (2006) Null mutations in progranulin cause ubiquitin-positive frontotemporal dementia linked to chromosome 17q21. Nature 442: 920–924. DOI 10.1038/nature05017

35. Hollingworth P, Harold D, Sims R, Gerrish A, Lambert JC, Carrasquillo MM, Abraham R, Hamshere ML, Pahwa JS, Moskvina V, et al. (2011) Common variants at ABCA7, MS4A6A/MS4A4E, EPHA1, CD33 and CD2AP are associated with Alzheimer’s disease. Nat Genet 43: 429–435. DOI 10.1038/ng.803

36. Naj AC, Jun G, Beecham GW, Wang LS, Vardarajan BN, Buros J, Gallins PJ, Buxbaum JD, Jarvik GP, Crane PK, et al. (2011) Common variants at MS4A4/MS4A6E, CD2AP, CD33 and EPHA1 are associated with late-onset Alzheimer’s disease. Nat Genet 43: 436–441. DOI 10.1038/ng.801

37. Sierksma A, Lu A, Mancuso R, Fattorelli N, Thrupp N, Salta E, Zoco J, Blum D, Buee L, De Strooper B, et al. (2020) Novel Alzheimer risk genes determine the microglia response to amyloid-beta but not to TAU pathology. EMBO Mol Med 12: e10606. DOI 10.15252/emmm.201910606

38. Bartels T, De Schepper S, Hong S (2020) Microglia modulate neurodegeneration in Alzheimer’s and Parkinson’s diseases. Science 370: 66–69. DOI 10.1126/science.abb8587

39. Paolicelli RC, Bolasco G, Pagani F, Maggi L, Scianni M, Panzanelli P, Giustetto M, Ferreira TA, Guiducci E, Dumas L, et al. (2011) Synaptic pruning by microglia is necessary for normal brain development. Science 333: 1456–1458. DOI 10.1126/science.1202529

40. Parkhurst CN, Yang G, Ninan I, Savas JN, Yates JR, 3rd, Lafaille JJ, Hempstead BL, Littman DR, Gan WB (2013) Microglia promote learning-dependent synapse formation through brain-derived neurotrophic factor. Cell 155: 1596–1609. DOI 10.1016/j.cell.2013.11.030

41. Nimmerjahn A, Kirchhoff F, Helmchen F (2005) Resting microglial cells are highly dynamic surveillants of brain parenchyma in vivo. Science 308: 1314–1318. DOI 10.1126/science.1110647

42. Davalos D, Grutzendler J, Yang G, Kim JV, Zuo Y, Jung S, Littman DR, Dustin ML, Gan WB (2005) ATP mediates rapid microglial response to local brain injury in vivo. Nat Neurosci 8: 752–758. DOI 10.1038/nn1472

43. Husemann J, Loike JD, Anankov R, Febbraio M, Silverstein SC (2002) Scavenger receptors in neurobiology and neuropathology: their role on microglia and other cells of the nervous system. Glia 40: 195–205. DOI 10.1002/glia.10148

44. Cheng C, Hu Z, Cao L, Peng C, He Y (2019) The scavenger receptor SCARA1 (CD204) recognizes dead cells through spectrin. J Biol Chem 294: 18881–18897. DOI 10.1074/jbc.RA119.010110

45. Hickman SE, Allison EK, El Khoury J (2008) Microglial dysfunction and defective beta-amyloid clearance pathways in aging Alzheimer’s disease mice. J Neurosci 28: 8354–8360. DOI 10.1523/JNEUROSCI.0616-08.2008

46. von Bernhardi R, Cornejo F, Parada GE, Eugenin J (2015) Role of TGFbeta signaling in the pathogenesis of Alzheimer’s disease. Front Cell Neurosci 9: 426. DOI 10.3389/fncel.2015.00426

